# Predicting natural language descriptions of smells

**DOI:** 10.1101/331470

**Authors:** E. Darío Gutiérrez, Amit Dhurandhar, Andreas Keller, Pablo Meyer, Guillermo A. Cecchi

## Abstract

There has been recent progress in predicting whether common verbal descriptors such as “fishy”, “floral” or “fruity” apply to the smell of odorous molecules. However, the number of descriptors for which such a prediction is possible to date is very small compared to the large number of descriptors that have been suggested for the profiling of smells. We show here that the use of natural language semantic representations on a small set of general olfactory perceptual descriptors allows for the accurate inference of perceptual ratings for mono-molecular odorants over a large and potentially *arbitrary set of descriptors*. This is a noteworthy approach given that the prevailing view is that human’s capacity to identify or characterize odors by name is poor [1, 2, 3, 4, 5]. Our methods, when combined with a molecule-to-ratings model using chemoinformatic features, also allow for the zero-shot learning inference [6, 7] of perceptual ratings for *arbitrary molecules*. We successfully applied our semantics-based approach to predict perceptual ratings with an accuracy higher than 0.5 for up to 70 olfactory perceptual descriptors in a well-known dataset, a ten-fold increase in the number of descriptors from previous attempts. Moreover we accurately predict paradigm odors of four common families of molecules with an AUC of up to 0.75. Our approach solves the need for the consuming task of handcrafting domain specific sets of descriptors in olfaction and collecting ratings for large numbers of descriptors and odorants [8, 9, 10, 11] while establishing that the semantic distance between descriptors defines the equivalent of an odorwheel.

---

To investigate whether semantic representations derived from language use could be applied to reliably predict ratings of a large set of detailed olfactory perceptual descriptors, we chose to predict the ratings of 146 fine-grained odor descriptors of the well known Dravnieks data set (see Fig.1a) [12]. We used as a starting point the ratings from 19 general descriptors in the DREAM data set [13] as it has 58 molecules in common, from 128 in total, and shares 10 descriptors with the Dravnieks data set (see Fig.1a and Extended Data Table 1). To quantify the semantic relationship between the DREAM and Dravnieks descriptors, we used a representation of linguistic data known as distributional semantic models. These models are quantitative, data-driven, vectorial representations of word meaning motivated by the distributional hypothesis, which asserts that the meaning of a word can be inferred as a function of the linguistic contexts in which it occurs [14]. A distributional semantic model assigns a vector to each word in a lexicon, based on the word’s use in language; words that are used in similar contexts, thus assumed to be more semantically similar, have vectors that are closer together in the distributional semantic space of the model (see Fig.1a bottom). In particular we utilized publicly available 300-dimensional semantic vectors produced using the fastText skip-gram algorithm that were trained on a corpus of over 16 billion words [15]. The fastText model contained vectors corresponding to the 19 DREAM descriptors which we refer to as the *DREAM semantic vectors*, and to 131 of the 146 Dravnieks descriptors which we refer to as the *Dravnieks semantic vectors*. Note that the training corpus was not biased in any way to include more or less olfaction-or perception-related material, i.e. it was intended to represent the general structure of semantic knowledge.

**Figure 1:**
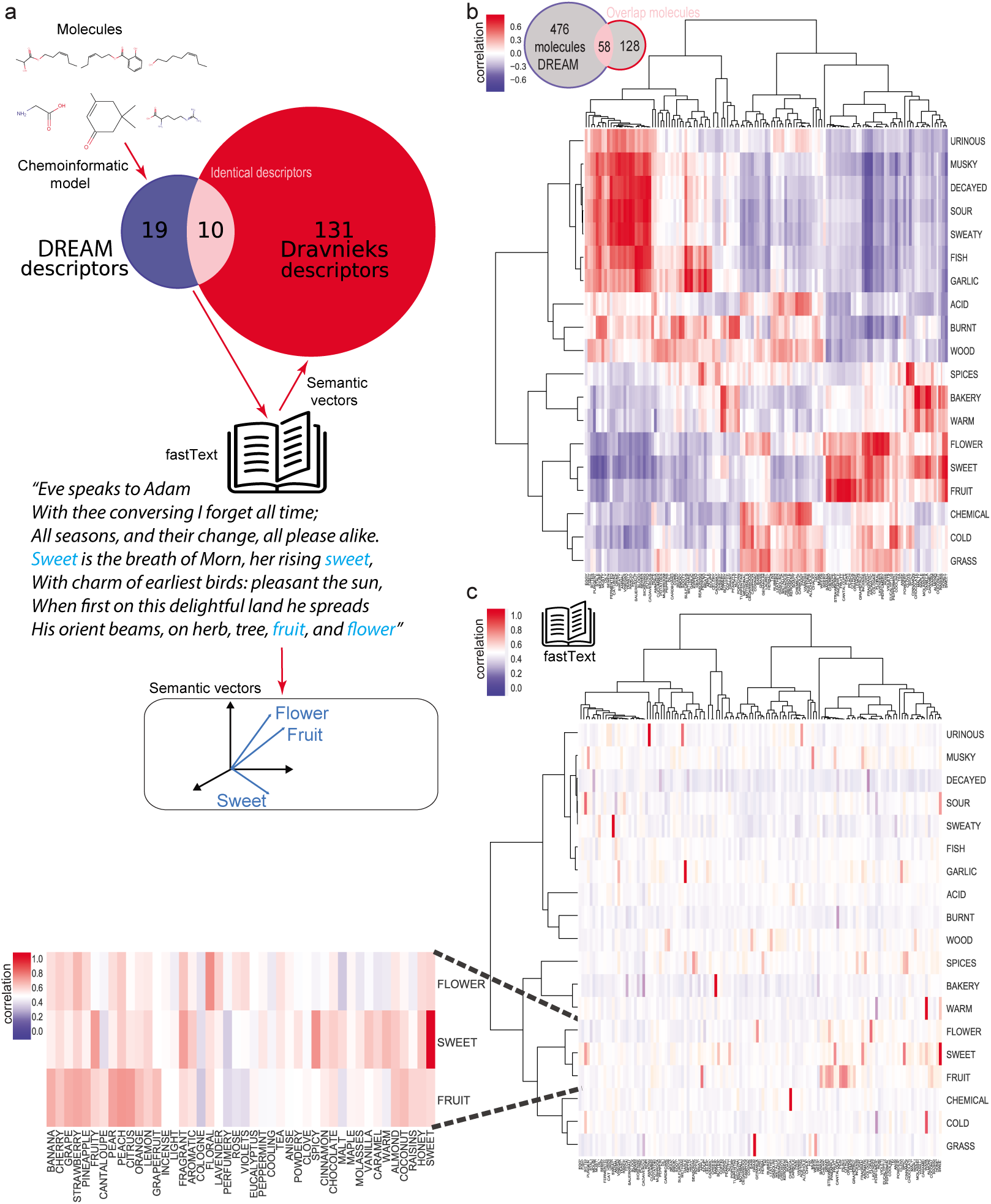
Construction of a universal perceptual map. **a.** Diagram showing the approach to predict rating for the Dravnieks descriptor sets. A chemoinformatic model helps predicting ratings for values of the DREAM set of 19 perceptual descriptors, the Dravnieks set has 146 descriptors, 10 descriptors overlap. We then use fastText to generate semantic vectors for the DREAM and Dravnieks descriptors by searching for co-occurrence of words in sentences as shown in the example (a fragment of Milton’s *Paradise Lost*). A model using these vectors is then applied to DREAM ratings to predict Dravnieks ratings for 131 descriptors. **b.** Heatmap showing correlations between the ratings of the Dravnieks descriptors (horizontal axis) and the DREAM descriptors (vertical axis) across the 58 shared molecules. The DREAM data set has ratings for 476 molecules, the Dravnieks data set for 128 molecules and they have 58 molecules are common to both data sets. Descriptors are arranged using hierarchical clustering, showing that they naturally cluster into semantically interpretable categories in the perceptual ratings correlation space. **c.** *Right*: Heatmap showing the correlation for the semantic vectors of the DREAM descriptors (horizontal axis) to the semantic vectors of the Dravnieks descriptors (vertical axis). Descriptors are arranged using the hierarchical clustering of **b** in order to allow direct comparison and emphasize common structure. *Left*: zooming in on one of the semantic clusters. Weights are not always highest between identical Dravnieks and DREAM descriptors (e.g., “flower”).

**Table 1:** Maximum correlation between DREAM (left) and Dravnieks (right) descriptors across 58 overlapping molecules.

|  |  |  |
| --- | --- | --- |
| BAKERY | MALTY | 0.870324290327 |
| SWEET | SWEET | 0.799501672662 |
| FRUIT | PEACH | 0.809634562228 |
| FISH | SAUERKRAUT | 0.841566522847 |
| GARLIC | SAUERKRAUT | 0.807089110708 |
| SPICES | CLOVE | 0.78750270191 |
| COLD | COOLING | 0.640995565798 |
| SOUR | RANCID | 0.85291315729 |
| BURNT | SMOKY | 0.611806648709 |
| ACID | CHEMICAL | 0.558275863815 |
| WARM | BAKERY | 0.629423235915 |
| MUSKY | SWEATY | 0.741410635391 |
| SWEATY | RANCID | 0.813893445052 |
| AMMONIA | URINE | 0.516659287626 |
| DECAYED | PUTRID | 0.796734051932 |
| WOOD | PEPPERS | 0.439374864874 |
| GRASS | HERBAL | 0.569829062717 |
| FLOWER | FLORAL | 0.779178189326 |
| CHEMICAL | CARBOLIC | 0.65387018244 |

Given that the DREAM and Dravnieks studies presented different sets of descriptors to the subjects, we expect that the perceptual ratings for the molecules in common will be re-anchored according to the available descriptors, and consequently that the descriptor ratings for the two datasets will differ even on shared descriptors [16]. Indeed we find that, although the correlations across the 58 shared molecules are high for the 19 corresponding descriptors (Fig.1b), the highest correlation is not always between the matching descriptors: e.g., although “sweet” in DREAM is most highly correlated to “sweet” in Dravnieks, “fruit” has a higher correlation to “peach” than to “fruity” (Extended Data Table 1). Nonetheless, the clusters of highly correlated descriptors defined by the dendrogram follow the close semantic relationship between the descriptors—e.g “flower” from DREAM correlates highly with the co-clustered “rose”, “violets”, “incense”, “perfumery”, “cologne”, “floral” and “lavender” from Dravnieks. We compared the correlation matrix based on the descriptors’ perceptual ratings (Fig.1b) to a correlation matrix between the DREAM and Dravnieks semantic vectors (Fig.1c right). We observe that the two correlation matrices are similarly structured (Procrustes dissimilarity *p <* 0.05 tested against randomized surrogates, correlation between maxima across the DREAM descriptors is *r* = 0.74, *p <* 10^−4^ and *r* = 0.5, *p <* 10^−9^ across Dravnieks descriptors). This is also reflected in the semantic vector correlation matrix where “sweet” is similarly maximally correlated with “sweet” in Dravnieks and “fruit” correlation is with “peach” and “citrus” than with “fruity”. Finally although “flower” shares a large weight with “floral”, it has similar correlation with “strawberry”, “fragrant”, and “lavender” (Fig.1c left).

The similarities in how the descriptors are arranged in the olfactory-perceptual space and the semantic space favor the hypothesis of a tight perceptual-linguistic bond between the descriptors ratings and their linguistic meanings. Consequently we developed a model that learns a transformation S from the 19 DREAM semantic vectors to the 131 Dravnieks semantic vectors (Fig.2a top left) and refer to this model as the direct semantic model. We hypothesized that, given the correspondences between the perceptual and semantic spaces, we could use S to predict the ratings of the 131 Dravnieks descriptors based solely on the ratings of the 19 DREAM descriptors and the semantic relation between the DREAM and Dravnieks descriptors. We compared the results of the semantic model to a direct ratings model that uses a training set of molecules for which both DREAM and Dravnieks ratings are available to learn a transformation R that can predict a new molecule’s ratings on the Dravnieks descriptors, given its ratings on the DREAM descriptors (Fig.2a top right). To further investigate the complementarity of the information provided by the semantic vectors and ratings data, we also looked at the performance of a mixed model that averaged the predictions of the two models.

**Figure 2.**
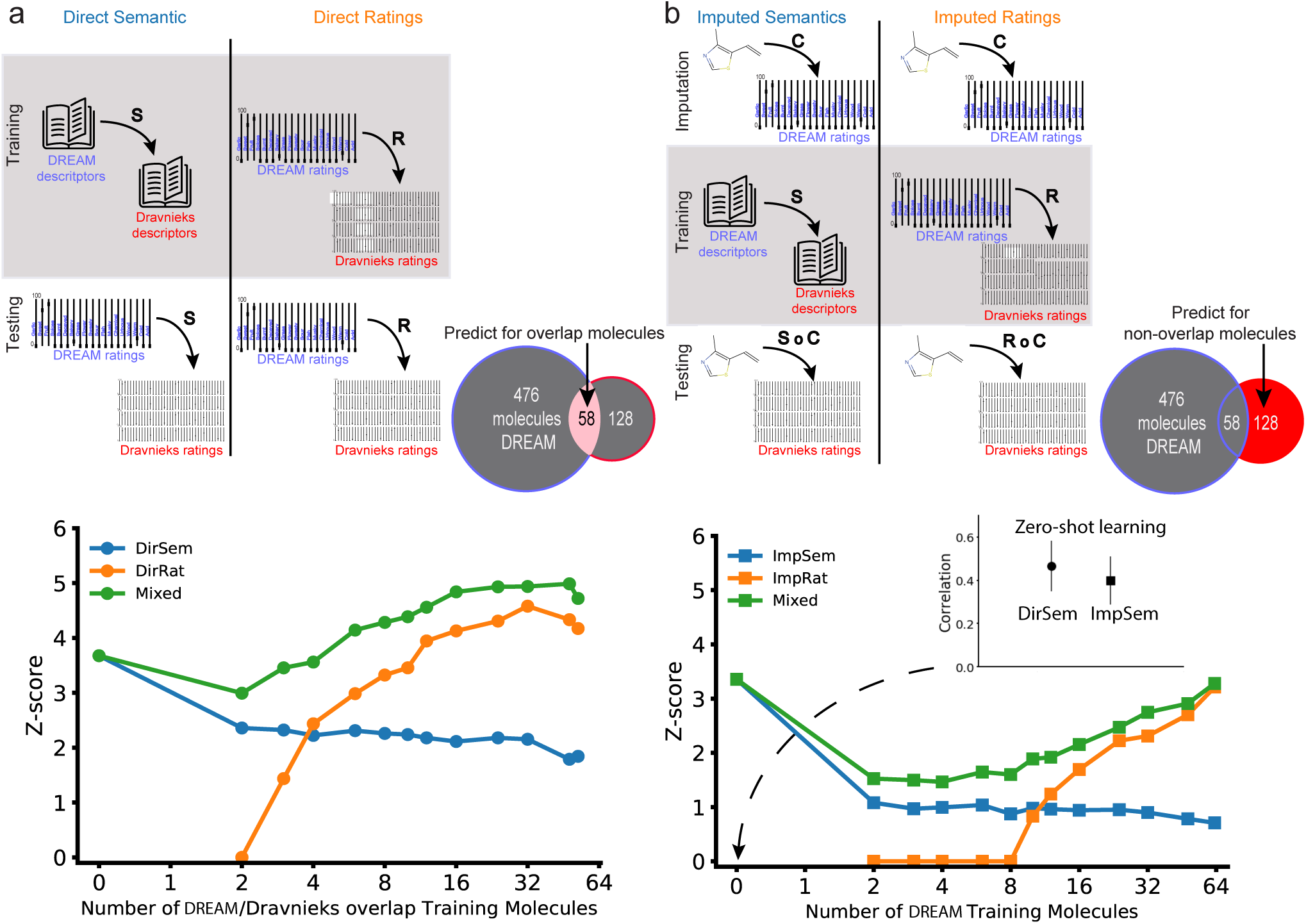
Predicting olfactory perception across descriptor sets and molecules. **a.** *Top* Schematic of the direct models for predicting ratings. During training (top row), the direct se-mantic model (*DirSem* left column) learns a transformation S from the DREAM descriptors’ corresponding semantic vectors to the Dravnieks descriptors’ semantic vectors. The direct ratings model (*DirRat* right column) learns a transformation R from ratings given to molecules on DREAM descriptors to ratings given to these molecules on Dravnieks descriptors. During testing (bottom row), the *DirSem* and *DirRat* models use the transformations S and R, respectively to predict the ratings given to molecules on Dravnieks descriptors from the ratings given these molecules on DREAM descriptors. Note that the *DirSem* model uses no molecules during training, while the *DirRat* model uses molecules from the shared set of 58 molecules during training. Both models are tested on these 58 molecules, averaging across 100 repetitions of 10-fold cross-validation. *Bottom*: The performance of the *DirSem* (blue dots) and *DirRat* (orange dots) models as well as a the averaged mixed (green dots), as the number of molecules used in training is increased. **b.** *Top*: Schematic of the indirect models for predicting ratings. During imputation (top row), both models learn the same transformation C from chemoinformatic properties to the ratings on the DREAM descriptors for molecules represented in the DREAM data set. During training (middle row), the two models imputed semantics *ImpSem* and imputed ratings *ImpRat* learn transformations S and R using the same procedure as the training phase of *DirSem* and *DirRat*, respectively. During testing (bottom row), the *DirSem* and *DirRat* models use the transformations S *˚* C and R *˚* C, respectively to predict the ratings given to molecules on Dravnieks descriptors from the ratings given these molecules on DREAM descriptors. Note that the *ImpSem* model uses no molecules during training, while the *ImpRat* model uses molecules from the set of 70 molecules present only in the Dravnieks dataset during training. Both models are tested on these 70 molecules, using cross-validation. *Bottom*: The performance of the *ImpSem* (blue squares) and *ImpRat* (orange squares) models as well as a the mixed model (green squares), as the number of molecules used in training is increased. *Inset* shows the value of the correlations for the *DirSem* (black dots) *ImpSem* (black squares) when no molecules are used during training, i.e. zero-shot learning.

To avoid overfitting, we used a cross-validation procedure where the 58 shared molecules are repeatedly divided at random into test sets and training sets and results averaged over repetitions. The performance of all three models was evaluated as the number of training molecules is varied. We compared each model’s performance by computing the median of the correlation between the predicted ratings and the actual ratings for a test set of molecules, across the Dravnieks descriptors. As ratings of molecules across descriptors are significantly correlated, we defined as an appropriate baseline prediction the mean rating for each descriptor across all molecules used for training the model and found that this baseline correlation is around 0.6. We then calculate a *Z*-score that compares the difference between the baseline correlation and the correlations produced by the models, taking into account their dependence. We report the median *Z*-score across molecules and across repetitions of cross-validation.

Remarkably, without making use of any of the ratings from the target set, i.e. an instance of zero-shot learning [6], the semantic model is able to predict the ratings in the target set reasonably well (Fig.2a bottom and 2b inset) with a median *Z* = 3.7, *r* = 0.47, *p <* 10^−4^ (see Extended Data Fig.1 for correlations plot) and better than the ratings model when trained on fewer than 6 overlapping molecules (Fig.2a bottom, blue and gold lines). Furthermore, the mixed model showed excellent performance with a *Z*-score of up to 5 and was never outperformed by the ratings model, underscoring the importance of the contribution from the semantic model and suggesting complementarity between information available in the ratings and the semantic model (Fig.2a bottom, green and gold lines).

We extended this approach using a chemoinformatics-to-perception model that allows the prediction of ratings along the 19 DREAM descriptors for any molecule using its molecular features [13]. We used an imputation model C, pre-trained with the DREAM dataset, to predict the 70 Dravnieks molecules that are not part of the DREAM dataset (Fig.2b top row; see also Methods). C is then combined with either the semantic transformation S to yield the imputed semantics model or used to train R yielding an imputed ratings model, both inferring Dravnieks ratings (Fig.2b middle row). These models were also averaged to produce a mixed model and scored on Dravnieks ratings (Fig.2b bottom row). Once again, predictions of descriptor ratings based on the semantic vectors alone with no molecular training data, are significantly better than chance when no training molecules are available (Fig.2b bottom, median *Z* = 3.4, *r* = 0.40, *p <* 0.001-see plot inset and Extended Data Fig.2 for correlations plot) and outperform the imputed ratings model when less than 10 molecules are available for training (Fig.2b bottom, blue and gold lines). We also again observe that a mixed model dominates the ratings model, showcasing the utility of semantic vectors even when ratings for a training set of molecules are available (Fig.2b bottom, gold and green lines). This advantage persists even as the number of molecules for which the source ratings available grows larger.

To understand the performance of the semantics-based models, we varied the number of source DREAM descriptors whose semantic vectors are available for training the direct and imputed semantic models while using leave-one-out cross-validation on their respective training/test molecule sets. The method we used for prioritizing the 19 perceptual descriptors is a state-of-the-art prototype selection method significantly superior to its competitors (see Methods) [17]. We observe that for both models, as the number of source descriptors increases, prediction performance generally increases, though the performance improvements plateau twice at four source descriptors and then around ten source descriptors (Fig.3a). The direct semantic model uses real DREAM ratings for making its predictions and so its correlation across descriptors is overall higher and the difference grows at the second plateau (Fig.3a squares and circles). This also suggests that it is possible to achieve good prediction performance on the target descriptors’ predictions by collecting only a small number of ratings from a smaller number of source descriptors.

**Figure 3.**
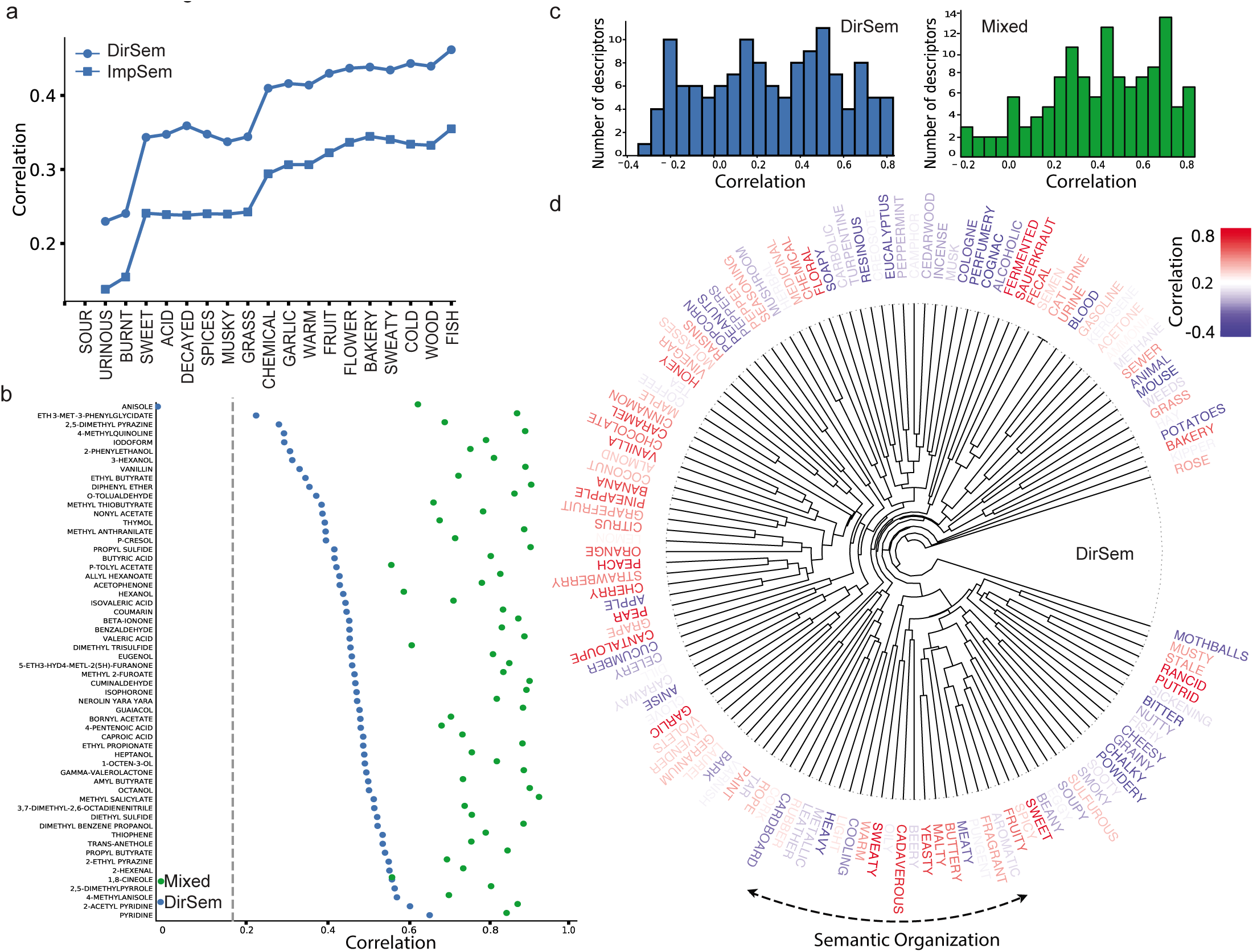
Analysis of predictive performance and map structure. **a.** The performance of the direct semantic *DirSem* and the imputed semantic *ImpSem* models (open blue circles and squares, respectively) as the number of descriptors used during training is increased. **b.** Pre-diction performance for each molecule, as measured by average correlation across descriptors between the ground truth ratings and the ratings predicted by the *DirSem* and mixed models (blue and green dots, respectively). The best-predicted molecules are toward the bottom of the chart, limit of significance (for *DirSem* model) is shown by the dotted gray line. **c.** Histograms showing the median correlation across molecules for each descriptor for *left* the *DirSem* model and *right* the mixed model. **d.** Odor wheel: prediction performance for each descriptor — as m asured by the correlation across molecules between the ground truth and the predictions from the *DirSem* model — is indicated by the color of the text (see bar for scale). Descriptors are arranged and clustered based on their semantic vectors.

**Figure 4.**
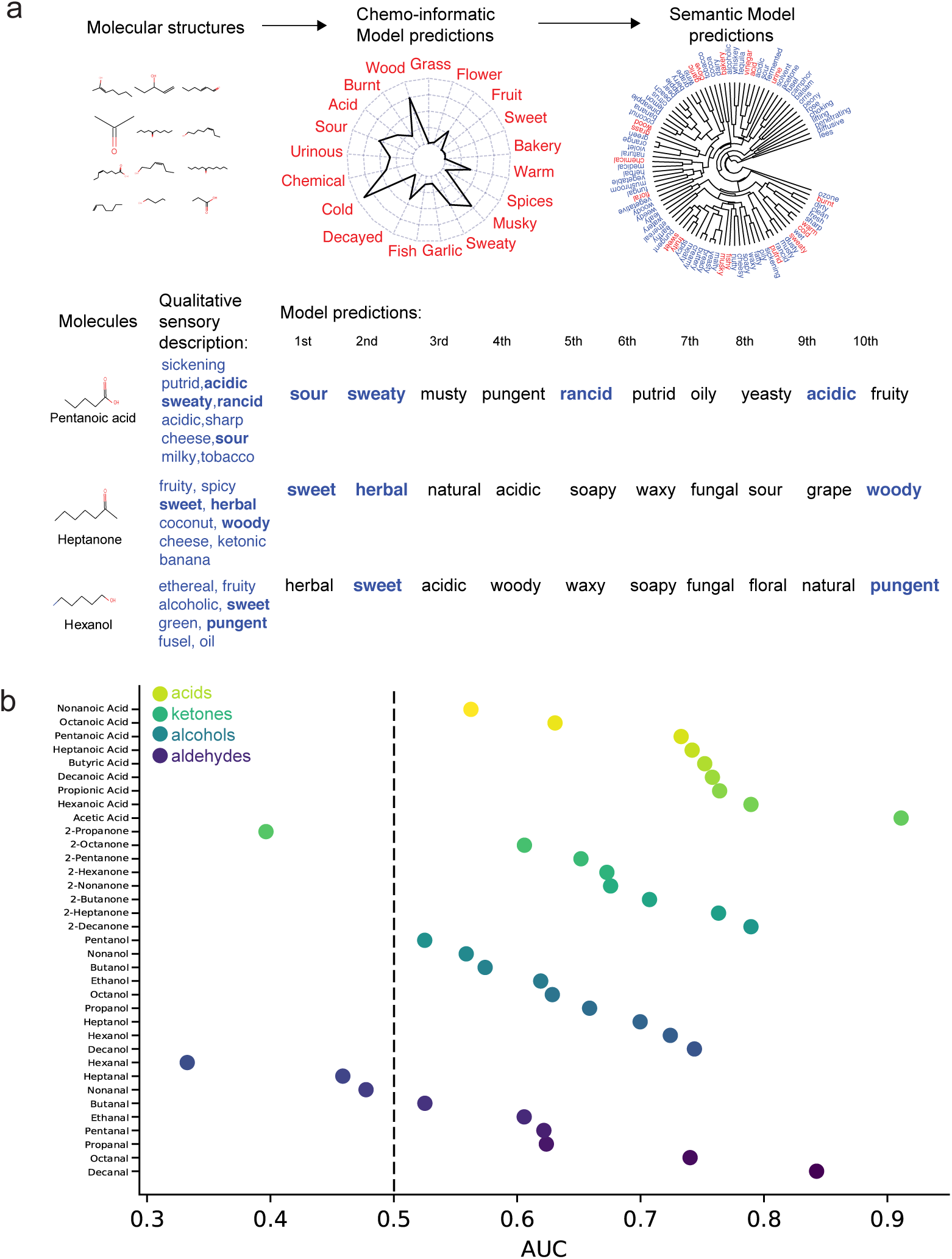
Predicting paradigm odors of four molecular families. **a.** Schematic for predicting paradigm odors for 35 molecules from the four chemical families of alkyl aldehydes, primary alcohols, 2-ketones, and carboxylic acids. textitTop. Features from molecular structures, *left*, are used to predict values of DREAM descriptors *middle*, and then the semantic model is applied to predict values in 80 unique perceptual descriptors extracted from the paradigm odor descriptions of all the 35 molecules, *right*. The 80 descriptors are shown in blue in a dendrogram according to their semantic similarity, the 19 DREAM descriptors are shown in red. The 7 overlap descriptors are acid, floral, fruity, sour, sweaty, sweet and wood. textit-Bottom. Example of performance of the model for 3 molecules *left*, their paradigm odors are indicated in blue *middle*, and predicted ordered list of 80 descriptors by decreasing ratings *right*, only first 10 are shown. Bold blue descriptors indicate a match to the paradigm descriptors of the molecule **b.** Prediction performance for the paradigm odors for each of the 35 molecules ordered by increasing AUC-ROC values for each of the four families of molecules starting with acids shown with dots in decreasing tones of yellow, ketones in decreasing tones of green, alcohols in increasing tones of light blue and aldehydes in increasing tones of dark blue. The dotted line indicates random AUC-ROC value.

We analyzed the quality of model predictions for each of the 58 overlapping molecules of this leave-one-out model (last green dot in Figure 2a and see Extended Data Table 2 for all the predictions) and find that the mixed model is more stable across molecules than the semantic model and as expected its correlations are also higher, around 0.8 on average (Fig.3b). Notably the semantic model predicted the perceptual ratings profile for 57 of the 58 shared molecules with significantly above-chance correlations. The best-predicted molecule *pyridine*, with a fish-like smell, had a correlation around 0.6 while the other top five predicted molecules had herbal and fruit-like smells (Fig.3b). We also analyzed the quality of the semantic model predictions for each of the 131 Dravnieks descriptors by displaying the median correlation across molecules for each descriptor in a histogram (Fig.3c left) and a dendrogram that accounts for their semantic organization (Fig.3d). Notably about 30 percent of descriptors were predicted with a correlation higher than 0.5 for the semantic model, a value that increased to 50 percent of the descriptors for the mixed model (Fig.3c right). Also, the prediction performance of the semantic model for a given descriptor is significantly correlated with the prediction performance of the nearest neighboring descriptor in semantic space (*r* = 0.4170, permutation test *p <* 0.001) and conversely a descriptor’s location in semantic space well predicts the prediction performance of the semantic model for that descriptor (*p <* .001 measured using 1- and 2-nearest-neighbors permutation tests). This smoothness in the prediction between descriptors reveals an odorwheel-like organization of the olfactory perceptual descriptors when organized by semantic distance.

To demonstrate the universality and flexibility of our zero-shot learning inference, we applied it to odor molecules that have been extensively studied by fragrance chemists and whose structure-odor relationship heuristics are well known. For this, we compiled notes on the smells of 35 molecules containing between two and ten carbon atoms in the homologous series of alkyl aldehydes, primary alcohols, 2-ketones, and carboxylic acids (see Methods) [18]. For each molecule, using the chemoinformatic and then semantic model method described above, we computed a prediction of the ratings for each of the 80 unique descriptors extracted from the smell notes (see Extended Data Table 3). We then ordered for each molecule the descriptors according to their ratings and computed the area-under-the-curve of the receiver-operating-characteristic curve (AUC) on the binary classification task of predicting whether the paradigm odors for each molecule contains the ordered descriptors (Fig.4). Notably, the family of acids were the best predicted family with a median AUC across molecules of 0.75 *p <* 0.02, ketones had an AUC of 0.67 and *p <* 0.05, alcohols had an AUC of 0.63 *p <* 0.07 and aldehydes were the worst predicted with an AUC of 0.61 *p <* 0.09. The overall median AUC across families of molecules was 0.66 with *p <* 0.05. Acids were overall predicted as sour but as the number of carbons increased the second-ranked descriptor changed from pungent to sweaty, musty and then back to pungent. Alcohols had overall an herbal smell and changed from sour to sweet (see Extended Data Table 3), aldehydes changed from pungent to sweet and fruity, finally 2-ketones changed from sour, acidic to sweet and grape. The limits of the semantic model are particularly clear for antonyms, generally side-by-side in the odorwheel. These limits are also reflected in the fact that synonymous odor descriptors have systematically different ranks. For example, each of the 35 molecules is predicted to be more “pungent” than “penetrating” with the median rank of “pungent” being 4 (our of 80) and the median rank of “penetrating” being 54 (out of 80).

There is a substantial body of evidence suggesting that the representations of words in semantic vector spaces obtained from co-occurrence statistics can be used to model human behavior [19, 20, 21, 22, 23, 24]. The present work demonstrates that the general structure of semantic knowledge, as manifested in the unbiased distribution of words in written language, can in fact be mapped onto the olfactory domain, creating a natural classification of olfactory descriptors, an odorwheel, that speaks to the depth of the connection between language and perception [25, 26, 27, 28, 29, 30].

This connection can be harnessed to effectively transform ratings from a small set of general descriptors to a larger more specific one. In combination with a chemoinformatics-to-perception model, our work enables end-to-end prediction of perceptual ratings for chemicals for which no ratings data is available at all, that is, a universal predictive map of olfactory perception. Given that specialists including tea and wine tasters, beer brewers, cuisine critics and perfumers expend considerable labor to set up lexicons that are concise and hierarchical, and which cover the relevant odor perception space, a general solution for predicting smell perceptual descriptors, independently of the lexicon used, would be highly impactful across a wide range of industries. Moreover, our findings are also clinically relevant, given that changes in olfactory perception are one of the first signatures of Alzheimer’s Disease [31] and associated with a range of other mental disorders [32]. Our approach provides a means to assess directly how these perceptual disturbances are associated with cognitive and emotional states.

**Online Content Methods**, Extended Data Figures 1-3 are available in the online version of the paper; references unique to these sections appear only in the online paper. Received –; accepted –. Published online –.

## Acknowledgments

We would like to thank Rick Gerkin and Pablo Polosecki for reading the manuscript and providing useful comments.

## Author Contributions

E.D.G developed the predictions and semantic model, A.D developed the chemoinformatic model. All authors together interpreted the results, approved the design of the figures and the text, which were prepared by E.D.G., G.A.C., and P.M.

## Author Information

Reprints and permissions information is available at www.nature.com/reprints. The authors declare no competing financial interests. Readers are welcome to comment on the online version of the paper. Correspondence and requests for materials should be addressed to P.M. and G.A.C.

## Methods

### Perceptual Data

In all of our experiments, we predict the average perceptual ratings given to molecules in the Dravnieks human olfaction data set [12]. This data set consists of the average ratings of 128 pure molecules by a total of 507 olfaction experts using 146 verbal descriptors. Each molecule was rated only by a subset of 100-150 of the experts. The ratings are on a scale from 0 to 5, where 5 signifies the best match of a descriptor for a given stimulus. Of these 146 descriptors, 15 were discarded because there was no corresponding word vector in our distributional semantic model (e.g., “burnt rubber”), leaving us with 131 descriptors.

Several of our models make use of the data collected by Keller and Vosshall [33] as presented in Keller et al. [13]. Data from 49 individuals were used, all of the work reported focuses on predicting the ratings averaged across subjects. Individuals were asked to rate each stimulus using 21 perceptual descriptors (intensity, pleasantness, and 19 descriptors), by moving an unlabeled slider. The default location of the slider was 0. The stimuli were 476 pure molecules. For each task, the final position of the slider was translated into a scale from 0 to 100, where 100 signifies the best match of a descriptor for a given stimulus. Further details on the psychophysical procedures and all raw data are available in the Keller and Vosshall article [33].

### Distributional semantic model

To assess accurately the semantic similarity between the DREAM and Dravnieks descriptors, we took advantage of a distributional semantic model trained using the fastText skip-gram algorithm, a neural network-based model that predicts word occurrence based on context [34]. These 300-dimensional vectors were trained on a corpus of 16 billion words, and are publicly available ^1^. See Bojanowski et al. [34] for additional details on training and the specifics of the model.

The semantic vectors of a distributional semantic model are vectorial representations of word meaning motivated by the distributional hypothesis stating that the meaning of a word can be inferred as a function of the linguistic contexts in which it occurs [14].

Distributional semantic models rest on the assumption that, to quote Wittgenstein, that ‘the meaning of a word is its use in the language’ [35]. For example, the distributional hypothesis would predict that *kitten* and *cat* have similar meanings, given that they are both used in contexts such as *the purred softly* and *the licked its paws*; meanwhile the meaning of *rock* would be less similar to *kitten*, because it is rarely if ever used in similar contexts. The distributional hypothesis has inspired the field of *distributional semantics*, which aims to quantify the meanings of words based on co-occurrence statistics of the words in large samples of written or spoken language. These co-occurrence statistics can be summarized and embedded in a low-dimensional vector space, known as a *semantic vector space*, using dimensionality reduction techniques such as principal components analysis [36] or neural networks [15]. The semantic vector space is constructed in such a way that words that occur in similar contexts and are therefore presumably semantically similar are represented by vectors that are geometrically close as measured for example by cosine distance or Euclidean distance.

### Chemoinformatic features

We used version six of the Dragon software package^2^ to generate a 4884 physicochemical features of each molecule (including atom types, functional groups, topological, and geometric properties)

### Estimating the perceptual ratings from chemical structure

To estimate the perceptual ratings from the chemical structure, we use a regularized linear model that is learned using elastic net regression [13]. This model is trained on the DREAM data set of 476 molecules. The input for the model consists of the chemoinformatic features of the molecules described above. Using these features, the model predicts the mean perceptual rating given by 49 subjects on each of the perceptual descriptors that we use above. Thus, for each molecule *i*, the chemoinformatics-to-perception model learns a transformation C such that

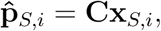

where 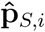 is the 19-dimensional vector containing the model’s estimate of the mean ratings on the DREAM descriptors for the molecule *i*, and x_*S,i*_ is the 4884-dimensional vector of molecule *i*’s chemoinformatic features.

### Extending ratings to new descriptor lexicons

We define two tasks, direct and imputed. For the direct task, we have access to actual DREAM ratings for each test molecule. In the imputed task, we do not have access to the test molecule’s actual DREAM ratings. Instead, we begin by applying a previously trained and unpublished model used in the context of Keller et al. [13] that can infer the ratings scores of any chemical on the DREAM verbal descriptors, given its chemoinformatic properties. For both tasks, the objective is to predict the test molecule’s Dravnieks ratings. Consequently, we also refer to the DREAM data as our *source* and the Dravnieks data as our *target*. We present three classes of model for each task, ratings, semantic, and mixed. Altogether the combination of the tasks and model classes results in six models, which we describe below.

As before, the real or imputed DREAM ratings scores for each molecule *i* can be collected into a 19-dimensional perceptual vector p_*S,i*_. In addition, for each DREAM descriptor *d*, we have a semantic vector s_*S,d*_, which is a 300-dimensional vector computed as described in the section describing of Word2vec. We collect these into a *source semantic matrix* S_*S*_ of dimension 19 *x* 300 where again 19 is the number of DREAM perceptual descriptors.

We want to learn the ratings scores for any arbitrary set of descriptors – we call these our *target* descriptors. We assume that we can compute the semantic vectors corresponding to each of these perceptual descriptors *d*, denoted by s_*T,d*_. Taking advantage of the structure inherent in these target semantic vectors is key to our method. We collect these into a *target semantic matrix* S_*T*_ of dimension *D*_*T*_ *x* 300 where *D*_*T*_ is the number of target descriptors. In the case of the results presented in the body of this paper, *D*_*T*_ = 131, because there are 131 Dravnieks descriptors that we use.

In this framework, our goal is to estimate the ratings scores for the target (Dravnieks) descriptors for each test molecule *i*, denoted by p_*T,i*_.

In order to set a point for comparison, we propose a baseline model that takes take the mean rating score for each target-set descriptor, across the training set of molecules for which ratings are available:

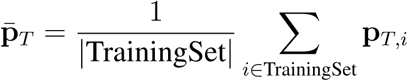

This is then used as the baseline estimate of the ratings scores across the target-set descriptors for a given new test molecule *i*. In the case where no training ratings are available for the target descriptors, we take the baseline to be the constant vector 0.

The first model class is composed of the semantics-only models for the direct and imputed tasks (*DirSem* and *ImpSem*, respectively). These semantics-only models assume that a distributional semantic space derived from a linguistic corpus shares structure with the olfactory perceptual space in which perceptual ratings scores exist. Consequently, we seek to test whether we can leverage the structure of the semantic space to predict ratings in the perceptual ratings space. To learn the semantics-only model S we proceed by supposing there exists a matrix S of dimension 19*x*131 that roughly maps from the semantic vectors for the source set of perceptual descriptors to the semantic vectors (collected into the matrix ∑_*S*_) for the target set of perceptual descriptors (collected into the matrix ∑_*T*_ :

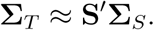

Our semantics-only models make the assumption that S is also an appropriate transformation for mapping from the perceptual ratings for the source set of descriptors to the perceptual ratings for the target set for each molecule *i*:

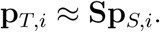

In order to estimate S, we use elastic net regression. The regularization parameters are set by nested 10-fold cross-validation.

Note that of the three model types described in this section, the semantics-only models are the only ones that do not rely on having access to any ratings scores for the source set (i.e., no p_*T*_ is required for training). However, to compare this model directly with the models that do use such information, we tested the effect of adding information about the mean rating to the model. Therefore, the final estimate for molecule *i* under this model, when target descriptors training molecules are available, would be:

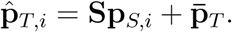

The only difference between *DirSem* and *ImpSem* is in the nature of p_*S,i*_ and 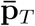. Recall that in *DirSem* these are derived from real DREAM ratings data, while in *ImpSem* they are predictions of the chemoinformatics-to-perception model.

The ratings-only models *DirRat* and *ImpRat* rely on having access to ratings scores for the target descriptors, for some training set of molecules. They assumes that there is some function R that maps from ratings scores on the source descriptors to ratings scores on the target descriptors for each molecule *i*:

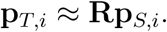

Once again, we estimate R using elastic net regression, with regularization weights set by nested 10-fold cross-validation. We also add information about the mean rating to the model, if available, so our final estimate under this model is:

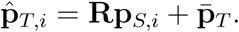

For the mixed models direct and imputed we simply average the predictions of the semantics-only and ratings-only models:

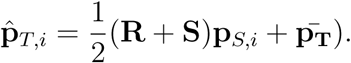

In preliminary investigations we also looked at other ways to combine the information in the semantics-only and ratings-only models, such as training a single regression model on the set union of the descriptors’ semantic vector values and molecule ratings, but a simple average performed best.

### Evaluating Performance

For each model, we vary the number of training molecules for which target descriptor ratings are available. We can then measure the median Pearson correlation between model *M* ‘s estimate 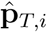 and the ground truth p_*T,i*_ for each test molecule *i* as:

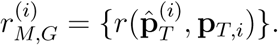

We use these correlations to assess whether the model’s performance differs significantly from the baseline model, by computing *Z*-scores. For the Semantics-Only model when we do not use any training molecules, the baseline is simply a correlation of zero, so the *Z*-score can be obtained using the Fisher *r*-to-*Z* transformation:

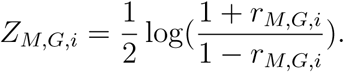

However, for the other models, note that the correlation coefficient produced by model and the correlation coefficient produced by the Baseline model are not independent random variables. Thus, to determine whether these two correlations differ significantly, we must take their dependence into account, which the standard Fisher transformation does not do. Instead, we can use the method developed by [37]:

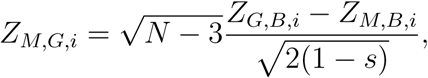

where

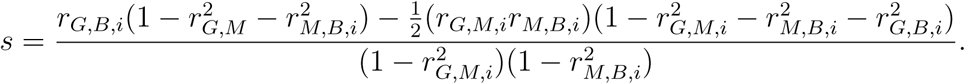

We can then compute the median of these *Z*-scores for all molecules in the test set:

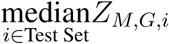

### Permutation tests for evaluating smoothness in semantic prediction

We performed a permutation test by randomly permuting the semantic nearest neighbors of each descriptor, and then re-computing the correlation between the prediction performances (measured by Pearson’s correlation) of each point and of its permuted nearest neighbor. The resulting simulated correlations exceeded the true correlation of *r* = 0.4170 in 0 of the 10,000 permutations.

For each descriptor, the *k*-nearest neighbor (*k*-NN) algorithm predicts the descriptor’s prediction performance (measured by Pearson’s *r*^2^) by taking the distance-weighted average of the prediction performance of the *k*-nearest neighbors. The mean squared error of this algorithm is then computed, and the significance is evaluated using a permutation test. The permutation test is performed by randomly permuting the semantic nearest neighbors of each descriptor, and then re-computing the mean squared error of the resulting *k*-NN predictions. The mean squared error of 2,000 such permutations was never below that of the true mean squared error.

### Tests for similarity between ratings and semantic vectors correlation matrices

To estimate the degree of structural similarity between the correlation matrix defined by Dravnieks and DREAM ratings (Fig. 1b), and that defined by the corresponding semantic vectors (Fig. 1c), we implemented two tests. In the first one, we computed the Procrustes dissimilarity between the rating matrix and the semantic matrix, and compared it against the expected dissimilarity between the original rating matrix and random permutation surrogates of the semantic matrix. A Wilcoxon test yields *p <* 0.05. For the second test, we found for each DREAM descriptor the Dravnieks descriptor with which it is maximally correlated, both in the ratings and semantic matrices. A Spearman test for the correlation between these two sequences yields *r* = 0.74, *p <* 10^−4^. Conversely, the test for the maxima estimated along the Dravnieks descriptors yields *r* = 0.5, *p <* 10^−9^.

### Additional information on elastic net regression

LASSO and elastic net are regression algorithms that impose a regularization penalty on the regression weights in order to reduce model complexity and avoid overfitting.

For a regression model of the form

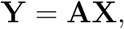

the regression weights in LASSO are estimated in order to minimize the following loss function:

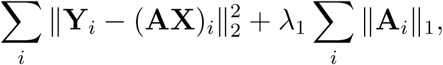

where the first term is the squared error of the prediction, and the second term is a regularization penalty (a penalty on the regression weights), and, *λ*_1_ is a regularization strength parameter. LASSO’s regularization penalty leads to a model that is sparse (i.e., produces few nonzero regression weights). This results in relatively more parsimonious and interpretable model. However, the LASSO loss function is not convex, so it does not produce a unique solution when the number of features is greater than the number of samples. When two features are highly correlated, LASSO will arbitrarily assign only one of the two features a nonzero weight, even if both contribute equally to the prediction in the ground truth model. This can lead to poor prediction performance.

Elastic net regression attempts to get around LASSO’s drawbacks. The regression weights are computed according to

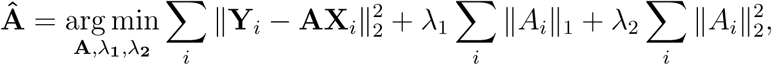

where the first term is the squared error of the prediction, the second term is the L1 (or LASSO) regularization penalty, the third term is the L2 (or ridge regression) regularization penalty [38], and, *λ*_1_ and, *λ*_2_ are the corresponding regularization strengths. Elastic net regression seeks to combine the benefits of LASSO and ridge regression. Like LASSO, it results in a parsimonious, interpretable, sparse model where most of the regression coefficients are zero. However, like ridge regression, elastic net has a convex loss function and produces a unique solution even when the number of features is greater than the number of samples. Elastic net also overcomes the arbitrary feature selection drawback of LASSO. See [38] for more details.

### Sequentially selecting prototypical features

We now describe the technical details of the method used to create Figure 3a. For a more thorough treatment please refer to [17].

Let *χ* be the space of all covariates from which we obtain the samples *X*^(1)^ and *X*^(2)^. Consider a kernel function *k* : *χ x χ →* ℝ and its associated reproducing kernel Hilbert space (RKHS) *К* endowed with the inner product *k*(x_*i*_, x_*j*_) = *〈 ∅* (x_*i*_), *∅* (x_*j*_) *〉* where *∅* **_x_**(y) = *k*(x, y) *∊ К* is continuous linear functional satisfying *∅* **_x_** : *h → h*(x) = *〈 ∅***_x_**, *h〉* for any function *h ∊ К* : *χ →* ℝ.

The maximum mean discrepancy (MMD) is a measure of difference between two distributions *p* and *q* where if ***μ****p* = 𝔼x∼*p* [*∅***_x_**] it is given by:

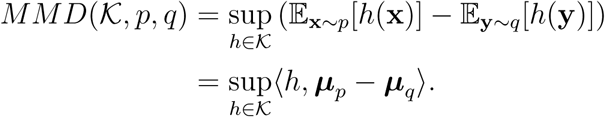

Our goal is to approximate ***μ***_*p*_ by a weighted combination of *m* sub-samples *Z ⊆ X*^(2)^ drawn from the distribution *q*, i.e., **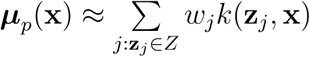** where *w*_*j*_ is the associated weight of the sample z_*j*_ *∊ X*^(2)^. We thus need to choose the prototype set *Z ⊆ X*^(2)^ of cardinality (*|.|*) *m* where *n*^(1)^ = *|X*^(1)^*|* and learn the weights *w*_*j*_ that minimizes the finite sample *MMD* metric with the additional *non-negativity constraint* for interpretability, as given below:

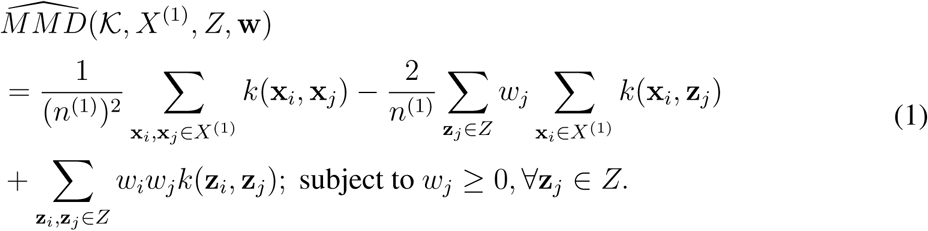

Index the elements in *X*^(2)^ from 1 to *n*^(2)^ = *|X*^(2)^*|* and for any *Z ⊆ X*^(2)^ let *L*_*Z*_ *⊆*[*n*^(2)^]= {1, 2, …, *n*^(2)^} be the set containing its indices. Discarding the constant terms in (1) that do not depend on *Z* and w we define the function

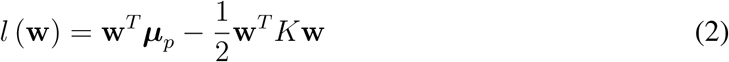

where *K*_*i,j*_ = *k*(y_*i*_, y_*j*_) and 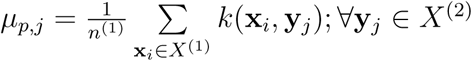 is the point-wise empirical evaluation of the mean ***μ***_*p*_. Our goal then is to find an index set *L*_*Z*_ with *|L*_*Z*_*| ≤ m* and a corresponding w such that the set function 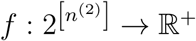 defined as

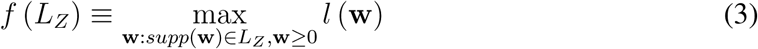

is maximized. Here *supp*(w) = {*j* : w_*j*_ *>* 0}. We will denote the maximizer for the set *L*_*Z*_ by 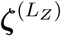.

The above problem is NP-hard to solve. The ProtoDash algorithm, however, efficiently solves this problem and is shown to have a tight approximation guarantee citeproto. If *Q* denotes the 476 x 19 [13] perceptual matrix then we set *X*^(1)^ = *X*^(2)^ = *Q*^*T*^ and run the following algorithm.

#### Algorithm 1 ProtoDash

**Input:** *X*^(1)^, *X*^(2)^

*L* = Ø ζ ^(*L*)^ = 0

g = ∇*l*(0) = ***μ*** _*p*_

*i* = 1

**while** *i* ≤ 19 **do**

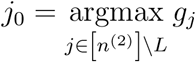

*L* = *L* ⋃ {*j*_0_}

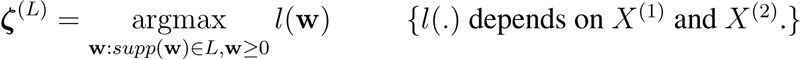

**g** = ∇*l*(ζ^(*L*)^) = *μ*_*p*_ − *K*ζ^(*L*)^

*i* = *i* + 1

**end while**

**return** *L*, *ζ* ^(*L*)^

The order in which elements are added to *L* is the order depicted in Figure 3a.

### Predictions of paradigm odors for molecular families

We extracted every term used to describe the paradigm odors for any of the 35 molecules in four families: 9 molecules from the family of alkyl aldehydes, 9 molecules from primary alcohols, 8 molecules from 2-ketones, and 9 molecules from carboxylic acids, that appeared in The Good Scents Company and Perfumer and Flavorist libraries. For these 35 molecules we included 80 descriptors used to describe all the molecules, ignoring instances where the term was only weakly associated – e.g. *fruity nuance* or *weak hint of apple*. This left us with a binary matrix of 35 molecules and 80 terms. We then proceeded to predict for these 35 molecules the 19 DREAM perceptual descriptors from the Dragon molecular descriptors of the molecules and then used the SEMANTIC model to obtain ratings for the 80 terms. Besides the AUC, we also computed for each molecule a *p*-value by performing a *t*-test for the difference between the means of the predictions for the terms that were used to describe the molecule and the terms that were not used to describe the molecule. A Kolmogorov-Smirnov test on these *p*-values reveals that they are not uniformly distributed (*p <* 1*e-*6), suggesting that overall predicted ratings for descriptors that are used to describe a molecule are ranked much higher than the predicted ratings for descriptors that are not used to describe a molecule.

Table 2: **Extended Data.** Predictions for leave-one-out models in Figure 3.

Table 3: **Extended Data.** Predictions for Paradigm Odors in Figure 4.

**Extended Figure 1.**
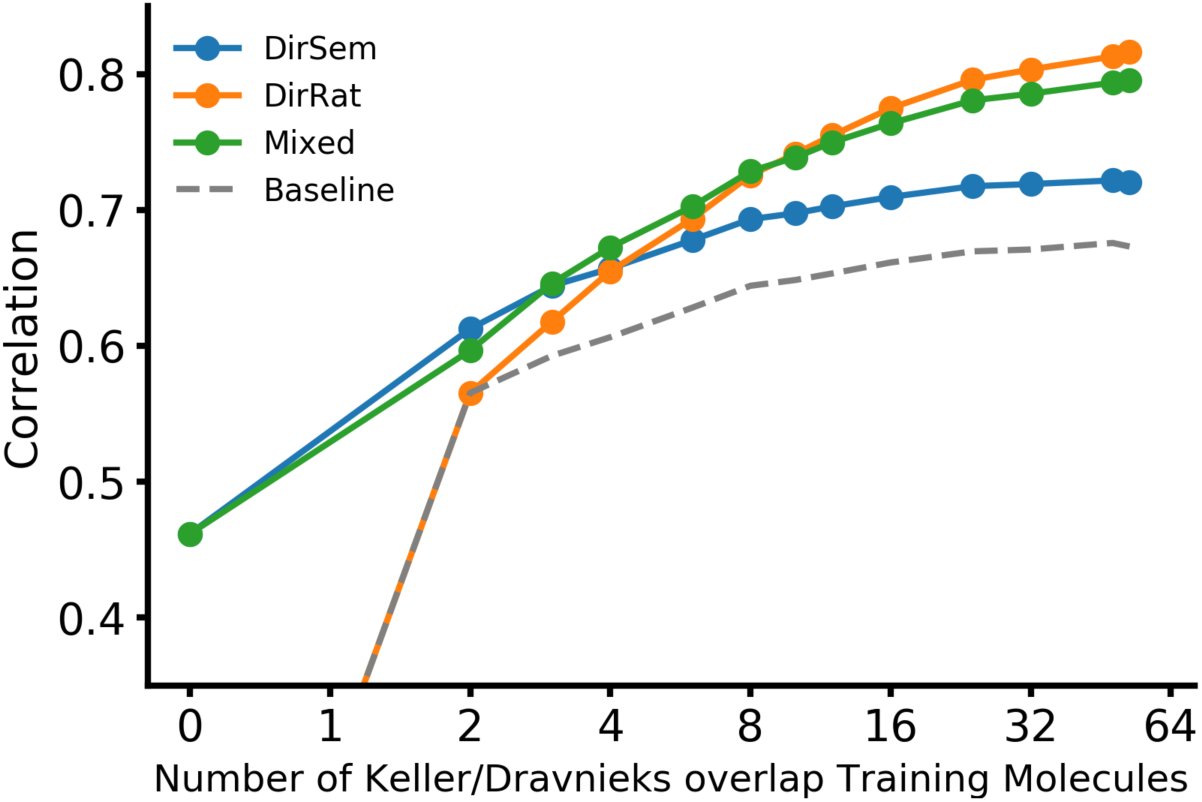
Maximum correlation for different models predicting Dravnieks descriptors across 58 overlapping molecules. The performance of the direct semantic (*DirSem* blue dots) and the direct ratings (*DirRat* orange dots) models as well as a the averaged mixed model (green dots), as the number of molecules used in training is increased.

**Extended Figure 2.**
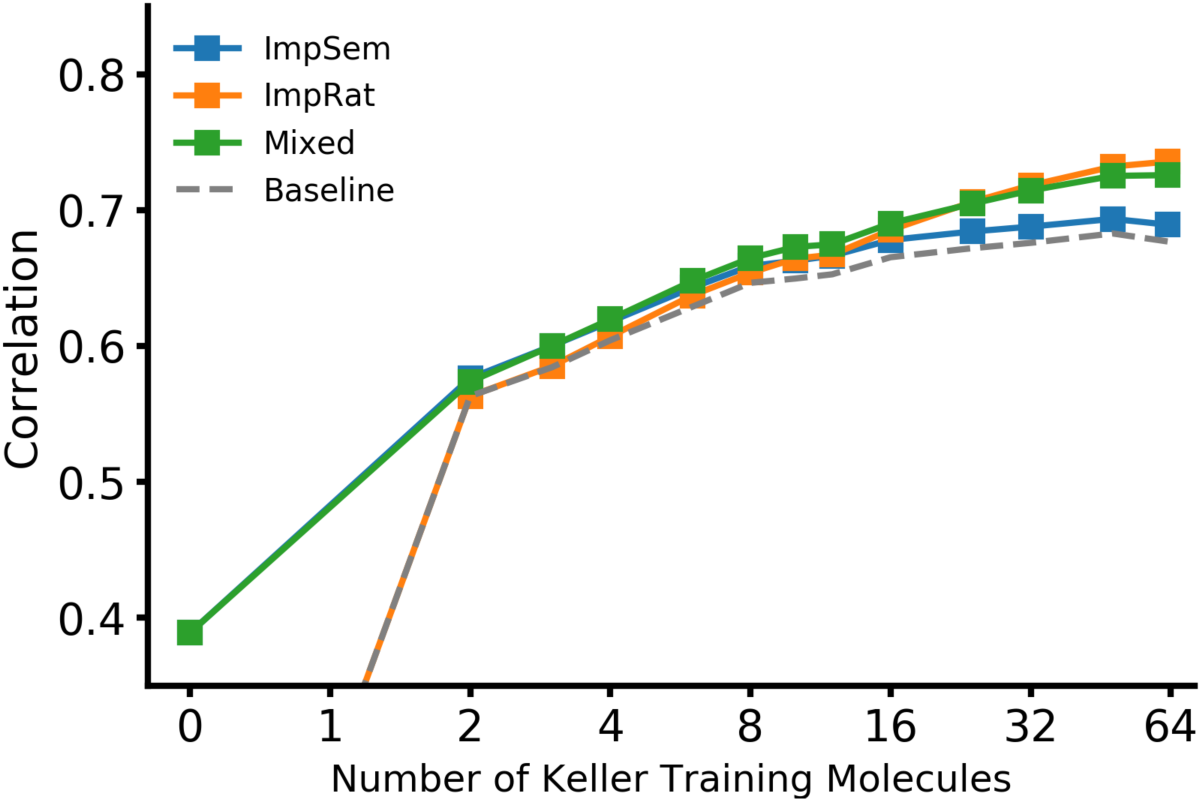
Maximum correlation for different models predicting Dravnieks descriptors across non-overlapping molecules. The performance of the imputed semantic (*ImpSem* blue squares) and the imputed ratings (*ImpRat* orange squares) models as well as a the averaged model (green squares), as the number of molecules used in training is increased.

## Footnotes

1 https://fasttext.cc/docs/en/english-vectors.html

2 http://www.talete.mi.it

